# Oncogenetic Network Estimation with Disjunctive Bayesian Networks

**DOI:** 10.1101/2020.04.13.040022

**Authors:** Phillip B. Nicol, Kevin R. Coombes, Courtney Deaver, Oksana Chkrebtii, Subhadeep Paul, Amanda E. Toland, Amir Asiaee

**Affiliations:** Harvard College, Cambridge, MA 02138, USA; Dept. of Biomedical Informatics, Ohio State University, Columbus, OH 43210; Natural Sciences Division, Pepperdine University, Malibu, CA 90263; Dept. of Statistics, Ohio State University, Columbus, OH 43210; Dept. of Cancer Biology and Genetics and Dept. of Internal Medicine, Division of Human Genetics, Comprehensive Cancer Center, Ohio State University, Columbus, OH, 43420; Mathematical Biosciences Institute, Ohio State University, Columbus, OH 43210

**Keywords:** cancer progression, Bayesian network, oncogenetic model, tumor phylogenetic

## Abstract

**Motivation:** Cancer is the process of accumulating genetic alterations that confer selective advantages to tumor cells. The order in which aberrations occur is not arbitrary, and inferring the order of events is challenging due to the lack of longitudinal samples from tumors. Moreover, a network model of oncogenesis should capture biological facts such as distinct progression trajectories of cancer subtypes and patterns of mutual exclusivity of alterations in the same pathways. In this paper, we present the Disjunctive Bayesian Network (DBN), a novel oncogenetic model with a phylogenetic interpretation. DBN is expressive enough to capture cancer subtypes’ trajectories and mutually exclusive relations between alterations from unstratified data.

**Results:** In cases where the number of studied alterations is small (*<* 30), we provide an efficient dynamic programming implementation of an exact structure learning method that finds a best DBN in the super-exponential search space of networks. In rare cases that the number of alterations is large, we provided an efficient genetic algorithm in our software package, OncoBN. Through numerous synthetic and real data experiments, we show OncoBN’s ability in inferring ground truth networks and recovering biologically meaningful progression networks.

**Availability:** OncoBN is implemented in R and is available at https://github.com/phillipnicol/OncoBN.

## 1. Introduction

Cancer is the process of accumulating molecular alterations that over time lead to cancer hallmarks (Hanahan and Weinberg, 2011). A natural question to ask is whether the order of alterations follows a particular pattern. Phylogenetic tree reconstruction methods answer this problem for individual tumors (Altrock et al., 2015). However, historically, due to the lack of high-resolution multi-region data of individual tumors, oncogenetic models were considered first. Oncogenetic models of tumorigenesis utilize many samples from the population of patients to estimate the order of alterations occur at the **disease** level, but are silent about the order of events at the **individual tumor** and **cell** levels. Recent technologies has enabled researchers to delineate various modes of evolution (Davis et al., 2017) and depict tumors’ evolutionary history in an unprecedented resolution (Gerstung et al., 2020). Although high-resolution data from individual tumors helps infer the tumor’s history, they do not provide the big picture of how a specific cancer type evolves. *In this work, we are taking first steps to reconciling these two levels of cancer progression modeling.*

The first oncogenetic model of tumorigenesis by Fearon and Vogelstein, 1990 was developed for colon cancer and suggested that a *chain* of aberrations is required to transform normal cells into carcinoma. Desper’s *Oncogenetic trees* (Desper et al., 1999) modeled progression as a rooted directed tree. Mixtures of oncogenetic trees (Beerenwinkel et al., 2005b,a) were proposed to capture the presence of an aberration in multiple progression paths. *Directed Acyclic Graphs* (**DAG**s) are the next straightforward generalization of tree-based models, as they allow multiple alterations (parents) to set up the clonal stage for the appearance of a new aberration (the child). *Bayesian networks* (**BN**), which are DAGs equipped with a joint probability distribution (Barber, 2012), lend themselves naturally to representing such models. Perhaps the most famous BN model of cancer progression is the *Conjunctive Bayesian Network* (**CBN**) (Beerenwinkel et al., 2007; Gerstung et al., 2009) which assumes all parent aberrations must be present in order for a child to occur.

The evolutionary interpretation of oncogenetic graphs is challenging. The most concrete biological way of thinking about an edge *e* = (*v, u*) in such DAGs is to assume mutation *v* fixates in the cell population and prepares the tumor for the next selective sweep by *u* (Gerstung et al., 2009). In other words, all mutations are assumed to be clonal, which is not accurate because of the observed intratumor heterogeneity in many cancer types (Dagogo-Jack and Shaw, 2018). *Our proposed tumorigenesis model has a phylogenetic interpretation and accommodates the presence of sub-clonal alterations.*

At its core, inferring cancer progression networks is the BN structure learning problem, which is NP-hard (Koller and Friedman, 2009). Various approximation and search algorithms have been proposed for cancer progression inference (Gerstung et al., 2009; Montazeri et al., 2016b; Farahani and Lagergren, 2013). These algorithms’ objective is to find a network structure that maximizes a (regularized) likelihood. The optimal network learned by any approximation method may be far from the ground truth and iterative search methods can get trapped in local maximums. *Here we show that for the number of driver alterations that we often encounter in tumors (< 30), one can use an efficient dynamic programming implementation of an exact structure learning algorithm* (Silander and Myllymäki, 2006).

### 1.1 Related Work

Mutual exclusivity of alterations is another phenomenon that was considered in learning cancer progression networks. Two sets of alterations are mutually exclusive if they (almost) never cooccur in a tumor (Leiserson et al., 2015). Two potential explanations for this observation are functional redundancy and synthetic lethality (Deng et al., 2019). Existing approaches considering pathways and their effects on cancer progression either assume that the pathways are inputs of the progression inference algorithm (Gerstung et al., 2011; Cheng et al., 2012) or learn them along with the progression network (Raphael and Vandin, 2015; Cristea et al., 2017).

The CBN progression rule dictates that all parent alterations need to be present in the tumor for the child to occur, under which mutually exclusive genes cannot share any descendant alterations. CBN’s inability to capture mutual exclusivity of alterations has motivated a line of work in which the mutual exclusivity restriction and pathway information are introduced artificially to the CBN (Gerstung et al., 2011; Cheng et al., 2012). Moreover, since each cancer subtype has distinct molecular characteristics and progression paths, one must first stratify samples to disjoint subtypes and then learn each subtype’s progression network separately. This extra step is required for all of the above models mainly because they cannot naturally capture subtypes’ mutual exclusivity. PICNIC (Caravagna et al., 2016) is the state-of-the-art pipeline that clusters samples to subtypes, detects driver events, checks for statistically significant mutual exclusivity hypotheses or takes pathway information as an input, and infers the progression network.

Several recent works attempt to model the accumulation of alterations by Suppes’ probability raising causal framework (Olde Loohuis et al., 2014; Ramazzotti et al., 2015; Caravagna et al., 2016; De Sano et al., 2016; Ramazzotti et al., 2018). Farahani and Lagergren, 2013 proposed (semi-)monotone progression networks without any biological interpretation. The class of montone BNs is a superset of our proposed model which makes it more flexible but prone to overfitting due to lack of enough samples in many real-world scenarios.

### 1.2. Our Contribution

#### Biological Modeling

We propose the Disjunctive Bayesian Network (**DBN**), which reconciles population-level progression models (oncogenetic models) and individual tumor evolution models (phylogenetic models). From the oncogenetic perspective, DBN relaxes the CBN progression assumption by allowing progression even if one of parents has occurred, Figure 1A. From the phylogenetic perspective, each directed path starting from the wild-type root in a DBN graph represents a (sub)clone, Figure 2C, and each sample from the DBN graph represents an individual tumor consisting of (sub)clones, Figure 2B. Overall, the DBN itself is the overlay of all of the possible sub-clones corresponding to the modeled cancer, Figure 2A. The DBN can naturally accommodate distinct progression paths for subtypes and is expressive enough to capture the mutual exclusivity of alterations present in the data. Therefore, one can skip two preprocessing steps necessary for the state-of-the-art models: stratifying samples by subtype and mutual exclusivity detection. We consider two extensions of DBN. The first extension relaxes the strict disjunction assumption and allows spontaneous (parent-less) alteration, Figure 1B. The second extension directly models measurement error of alterations. To have an uncluttered presentation, we present the measurement error model only in the Supplement A.

**Figure 1:**
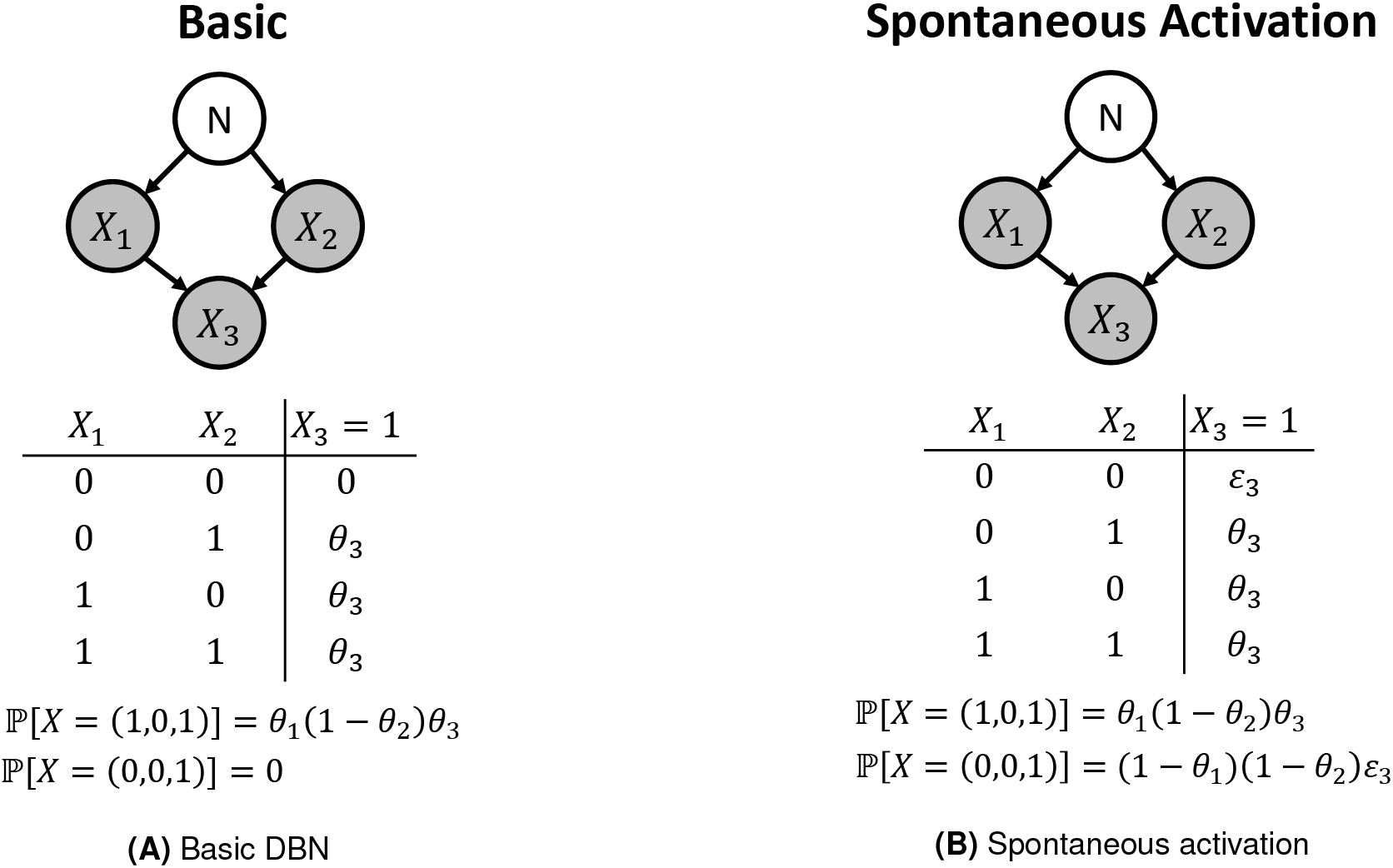
Bayesian networks of the cancer progression models investigated. Node *N* represents normal cell state, and each random variable *X_j_* is an observed alteration, and the corresponding progression probability parameter is *θ_j_*. In all models, the conditional probability table of *X*3 is shown, and probabilities of instance observations are computed. **(A)** Basic DBN model where further progression is impossible if none of the parent alterations have occurred. **(B)** Spontaneous activation model where there is a non-zero chance of a child occurring even if none of its parents are active.

**Figure 2:**
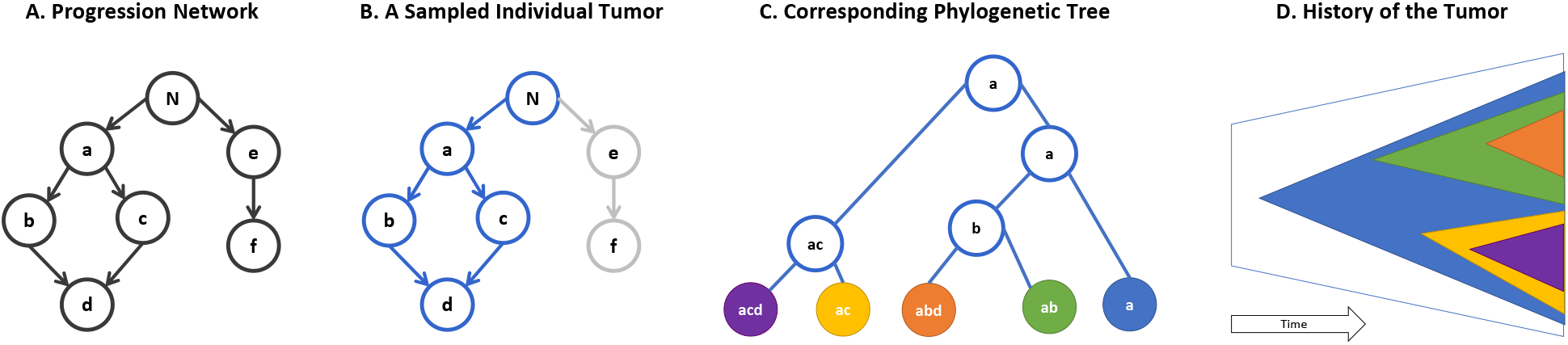
Phylogenetic interpretation of the DBN model. **A.** A DBN progression network that models a cancer type at the population level (disease level). Root *N* represents the wild type state (Normal) and there are six known driver alterations. **B.** A sample from the network (blue nodes) that represents an individual tumor. **C.** The corresponding phylogenetic tree of the sample. Each path of sampled graph forms a subclone living on the leaves of the phylogenetic tree that are distinguished by various colors. **D.** Visualization of the tumor history and the subclonal relationships through time.

#### Computational Efficiency

We provide an efficient Dynamic Programming (**DP**) implementation of an exact structure learning method (Silander and Myllymäki, 2006) that learns the optimal DBN (in terms of a regularized likelihood). Additionally, this algorithm can be incorporated into existing cancer progression frameworks such as Conjunctive Bayesian Networks (Gerstung et al., 2009) or CAPRI (Ramazzotti et al., 2015), which will likely improve their accuracies. For rare cases that the studied driver alterations are numerous, we provided an efficient Genetic Algorithm (**GA**) in our software package. To speed up the GA’s global search, we characterize a likelihood-equivalence relation over DBNs and only search through the representative DAGs of each class.

#### Experimental Performance

Through numerous synthetic and real data experiments, we show the ability of our algorithms in reconstructing ground truth progression networks from simulated samples and inferring biologically interpretable progression networks for cutaneous melanoma, lung adenocarcinoma, and bladder cancer. Our scalable **Onco**genetic **B**ayesian **N**etwork R package, **OncoBN**, provides two easy to use routines (approximate and exact) for estimation of onco-genetic Bayesian networks including DBN and CBN.

## 2. Methods

We model the observation of alterations as a binary random vector (*X*_1_,…, *X_p_*), where *X_j_* = 1 if the *j*-th alteration is detected in the sample and **x** = (*x*_1_,…, *x_p_*) is an observed sample. We assume that a BN governs the order in which the events can occur. The BN consists of a DAG *G* and local *Conditional Probability Distributions* (**CPD**) 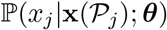 where 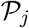 is the set of parents of event *j* in *G* and *θ* parameterizes the distribution. Local CPDs form the joint distribution as 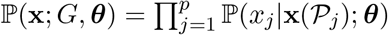.

### 2.1. Progression Rule and Parameter Estimation

#### Basic DBN

The DBN progression rule asserts that an event *j* occurs with probability *θ_j_* if and only if at least one of its parents have occurred. Therefore, 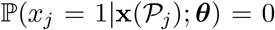 if parents are inactive and *θ_j_* otherwise, Figure 1A.

#### Spontaneous Activation Model

The deviation from the DBN progression rule may be the results of *spontaneous activation* caused by unknown sources. To capture that, we add a non-zero spontaneous activation probability *ε_j_* > 0 for each node, Figure 1B.

Given *n* cross-sectional samples and the network *G*, we wish to find 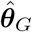, the maximum likeli-hood estimator (MLE) for ***θ***. We focus on the spontaneous activation model, where the likelihood is:

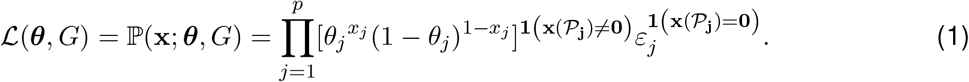

Maximizing the log-likelihood results in 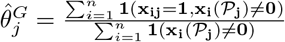 and where *x_ij_* is the realization of the *j*th event in the *i*th sample. From now on, to reduce the number of inferred parameters, we assume *∀j*: *ε_j_* = *ε* and we fix it throughout the experiments. Details of parameter estimation for the three models (basic, spontaneous, measurement error) are presented in Supplement B.

### 2.2. Exact Structure Learning

Although for a fixed network *G* the MLE parameters have closed form, finding the best *G* is NP-hard. We present an efficient Dynamic Programming (DP) method for *p <* 30 that finds a best graph with maximum likelihood. We use “a best” instead of “the best” graph to emphasize on the fact that the graph with the maximum likelihood is not unique.

To have more interpretability and avoid overfitting, we restrict our search space to the space of *p*-node DAGs with an in-degree bound of *k*, 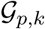. To further penalize dense graphs, we follow (Ramazzotti et al., 2015) and use the Bayesian information criterion (BIC) as our graph fitness score. The final optimization objective takes the following form:

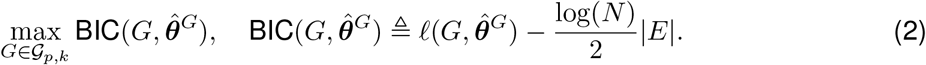

#### 2.2.1 Dynamic Programming Algorithm

An exhaustive search of 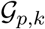 takes super-exponential time. Silander and Myllymäki, 2006 introduced a dynamic programming algorithm that can find the optimal network in exponential time. Their algorithm assumes that each graph *G* can be assigned a decomposable score Score(*G*) such that 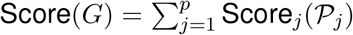 where 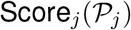 is the score of the subgraph consisting of only vertex *j* and its parents 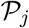. 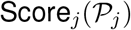 is called the local score of *j*. For us, 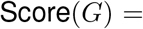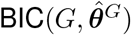, is our decomposable score. The rest of this section is devoted to a high-level summary of the algorithm.

##### Optimal Substructure

First note that each DAG has at least one sink node, which is a node with no outgoing edges. The score of a best graph *G*^*^(*V*) can be broken down to the best parents of any of its sinks *s* and a best subgraph obtained by removing *s* and its incoming edges. More formally, for *s*, an arbitrary sink of *G*^*^, 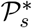 should be a best set of parents, i.e., has highest local score 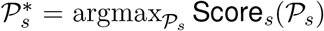. In addition, for *G*^*^(*V*) to be optimal, Score(*G*^*^(*V \{s}*)) should also be optimal. This optimal substructure suggests the following recursive formula for finding a best sink for set of nodes *W ⊆ V*:

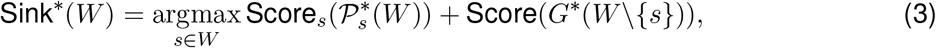

where 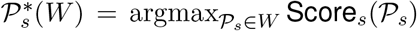 is the pre-computed best parents of *s* in *W*. Best sinks can be computed in *O*(*n*2^*n−*1^) time using memoization.

##### Reconstructing an Optimal Solution

Best sinks immediately result in a best ordering of nodes in reverse order. By having an optimal order and the best set of parents for all nodes, it is straight-forward to build an optimal graph. Starting from an empty graph, we add a node according to the optimal order and add incoming edges from its optimal parents that preexist in the graph.

##### Computational Complexity

The most intensive portion of the algorithm is computing the set of best parents 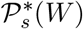 for every *W ⊆ V \{s}*. This step requires *O*(*n*^2^2^*n−*1^) time and *O*(*n*2^*n−*1^) space. By leveraging disk space, it is possible to implement the algorithm such that at most 2^*n*+2^ bytes of RAM are occupied at any given time.

#### 2.2.2. Pruning Spurious Edges

When the data is corrupted by noise, the estimated graph is likely to contain spurious edges. To remove low confidence edges, we perform statistical tests on the estimated graph. In DBNs, if *e* = (*u, v*) is an edge in the ground-truth graph, we have ℙ (*X_u_* = 1 | *X_v_* = 1) *>* ℙ (*X_u_* = 1 | *X_v_* = 0). Thus, we use the Fisher’s exact test to check the inequality and retain edges for which the inequality holds with high confidence.

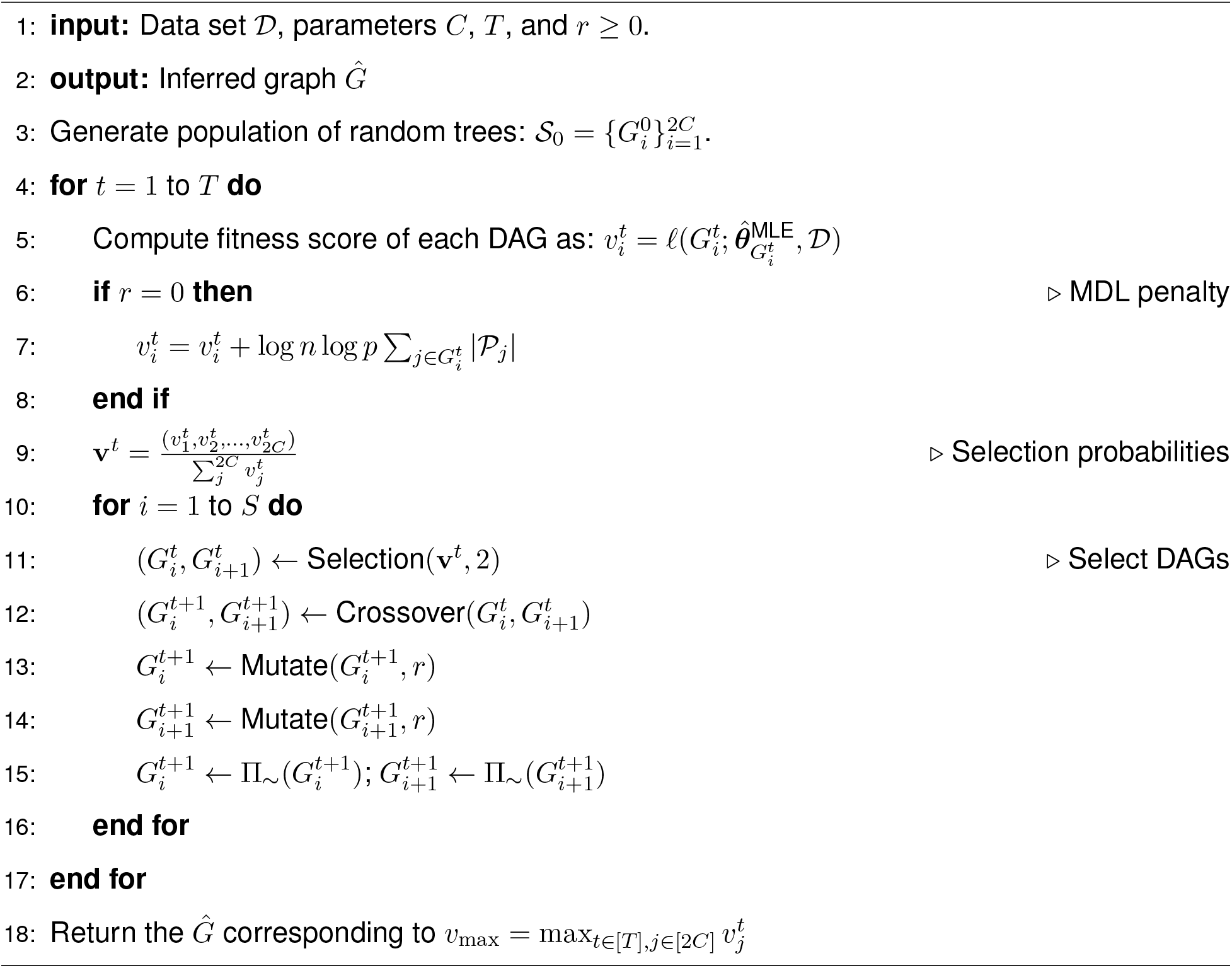
Algorithm 1. Genetic Algorithm of OncoBN Package

### 2.3. Approximate Structure Learning

For large number of mutations the exhaustive search is infeasible. Here we propose a Genetic Algorithm (**GA**) to approximate the global maximum to the log-likelihood function *l* for *p >* 30. The pseudocode of this part is summarized in Algorithm 1.

#### 2.3.1. Genetic Algorithm

Genetic algorithms searches for a global optimum using a “survival of the fittest” strategy. We begin with a population of 2*C* candidate solutions known as *chromosomes* and evolve them for *T generations*. Each chromosome is assigned a *fitness value v* which determines its quality. Then, *S* chromosome pairs are selected preferentially according to their fitness for reproduction. The next generation forms by performing a *crossover operation* on chromosome pairs. In each generation, there is a chance that a *mutation operation* changes chromosomes. In the setting of our model, chromosomes at generation *t* are 2*C* DAGs, 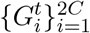 and the fitness of each DAG is its maximum likelihood value.

##### Representation

The most natural way to encode a DAG *G* is by using its adjacency matrix **A**. However, perturbing the entries in **A** may unintentionally introduce directed cycles into the resulting graph. To avoid this problem, following Carvalho, 2013, we represent *G* with a pair (**O**, **π**), where **O** is the adjacency matrix for the *topological ordering* of *G* (i.e., an strictly upper triangular matrix), and **π** is a permutation vector describing how the vertices of **O** should be relabeled to generate **A**, Figure 3. We consider the ordering **O** and permutation **π** as separate chromosomes and evolve each of them individually. We can avoid introducing directed cycles by ensuring that our genetic operators always return an upper triangular matrix.

**Figure 3:**
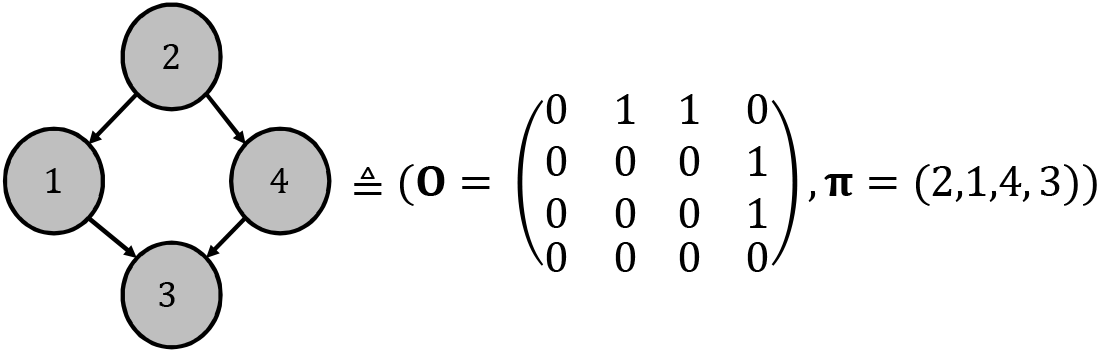
DAG representation. The DAG can be decomposed into an upper triangular matrix **O** along with a permutation ***π***.

##### Operations

Each crossover operation is defined to take in two DAGs and produce two offspring to keep the generation size constant. The orderings and permutations are crossed over separately (Supplement C). To maintain diversity in the population, we also define three mutation operators: edge, branch, permutation (Supplement C).

#### 2.3.2. Speeding up the GA with DAG Equivalence Classes

Since mutation *i* activates with probability *θ_i_ irrespective* of which parent mutations are active, many different network structures induce the same probability distribution over {0, 1}^*p*^. We say that *G ~ G*’ if, for every ***θ*** and **x**, ℙ(**x**; *G*, ***θ***) = ℙ(**x**; *G*’, ***θ***). It is clear that *~* defines an equivalence relation over DAGs. To make the GA more efficient, we search only one DAG per equivalence class by defining a *canonical form* for each graph. Figure 4 gives an example of equivalent networks. Algorithmically, we project back new solution graphs to the state space of canonical forms by removing redundant edges and uniquely labeling similar vertices in function Π_~_(·) (line 12 of Algorithm 1.) More details on mathematical properties of DBNs is presented in the Supplement D.

**Figure 4:**
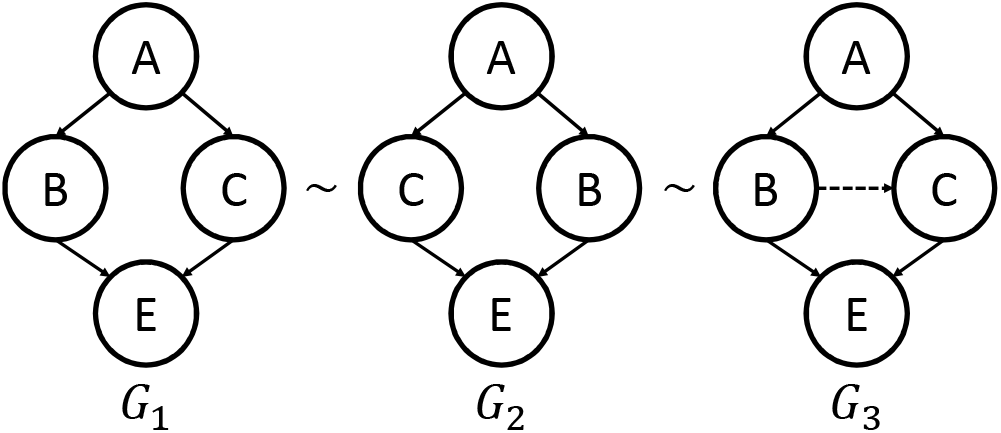
Examples of DAGs from the same equivalence class and their canonical form. For all ***θ*** and **x**, ℙ(**x**; ***θ***) is the same for all of the three network structures shown above. *B* and *C* are similar vertices in *G*_1_ and *G*_2_. Edge *B → C* is redundant in *G*_3_. By uniquely labeling similar vertices and removing redundant edges we reach *G*_1_ as the canonical form of the other two DAGs.

#### 2.3.3. Controlling Complexity

To prevent overfitting, we consider two types of penalty to control the complexity of the learned BN. First, if *r* = 0 in Algorithm 1, we perform regularized MLE by using the Minimum Description Length penalty introduced in (Lam and Bacchus, 1994) that simplifies to 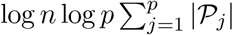 for the DBN. In another approach represented by *r >* 0 in Algorithm 1, we limit the number of parents of each node to *r*, i.e., 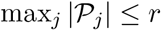.

## 3. Results

### 3.1. Inferring Simulated Ground Truths

To test the DP method against existing cancer progression algorithms, we generate datasets from simulated networks. Random graphs *G* are created using the PCALG R package (Kalisch et al., 2012), which allows the user to specify the number of vertices and the average degree. For network parameters, we sample *θ_j_ ~* Unif(0.25, 0.75). Once *θ_j_*s and *G* are known, a simulated dataset can be created by iterating over a topological sort of *G*. For tests on simulated data, we fix the number of observations *n* to be 400 and the number of alterations *p* to be 20 (this is similar to the size of existing cancer datasets). Unless specified otherwise, the average degree is set to 3. To simulate the noise that is likely present in real data, we flip the binary value of each entry with probability *η*.

If 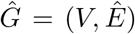 is the estimated network with ground truth *G* = (*V, E*), one can define a false positive edge to be an edge 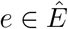 with *e ∉ E* and false negative edges similarly. Since the number of possible false positives is likely much larger than the number of possible false negatives, we assess performance using Matthew’s correlation coefficient (MCC), which is robust under uneven class sizes (Matthews, 1975). The MCC can be computed as

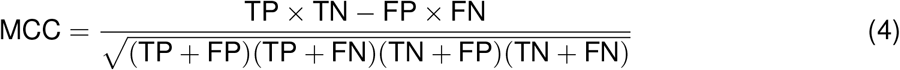

where TP (FP) is the number of true (false) positives and TN (FN) is the number of true (false) negatives. A MCC of 1 corresponds to perfect reconstruction, While an MCC of 0 means the algorithm is outputting a random network.

The DP algorithm requires that the spontaneous activation rate *ε* and in-degree bound *k* are chosen in advance. We suggest (and use) the following heuristic to set *ε*: set *ε* = *f_m_*/2, where *f_m_* is the frequency of the *least* frequent alteration. One should always select *ε < f_m_*, as otherwise there may be incentive to misplace the node corresponding to this alteration. In the interest of efficiency, we set *k* = 5, although in theory one could test every possible *k* to select the one that best trades expressivity for complexity. For pruning spurious edges, Fisher’s exact test with significance level of 10^*−*5^ is used.

First, we compare the DP algorithm to CBN. The original approach of Gerstung et al. (2009) uses simulated annealing to approximate the network structure alongside a computationally expensive expectation-maximization (EM) algorithm for parameter estimation. As a result, their method is only applicable when the number of mutations is less than 12. Montazeri et al. (2016a) addresses this issue by developing an efficient Monte Carlo algorithm, named MC-CBN, to estimate the parameters and structure of a CBN. Figure 5A compares MC-CBN and the DP algorithm for various choices of *η* ∈ [0, 0.2]. In the case of low error (*η ≈* 0) both methods are extremely accurate. However, as *η* becomes larger, the MCC for MC-CBN drops to 0 at a faster rate.

**Figure 5:**
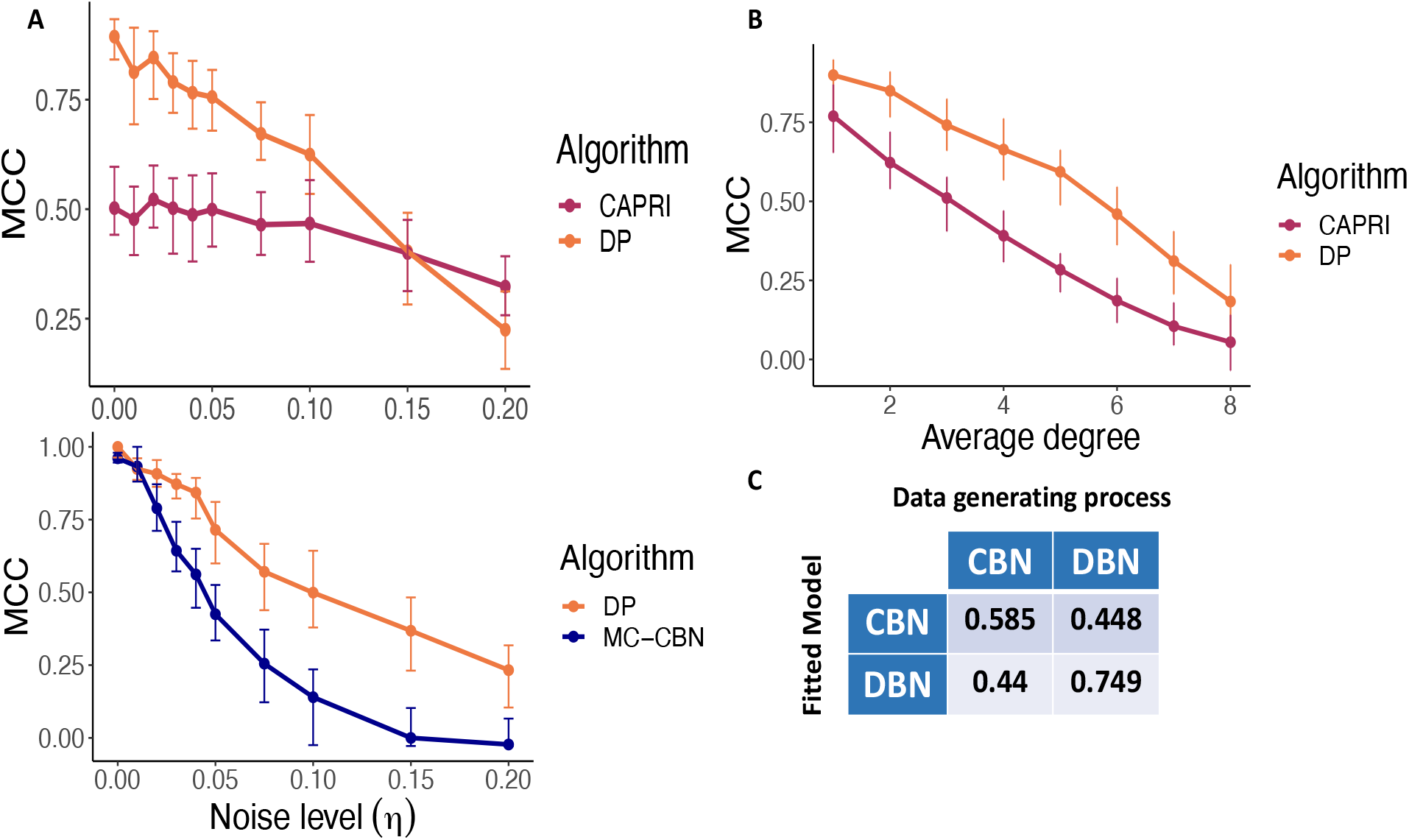
Comparison to existing cancer progression algorithms on simulated data. **A.** Comparing the DP algorithm to CAPRI and MC-CBN for various noise rates. For each value of *η*, 100 datasets were generated. Points represent the median MCC over all trials and error bars give interquartile range. **B.** Comparing the DP algorithm to CAPRI for various levels of network complexity. Network complexity was quantified by varying the average degree of random graphs from 1 to 8. 100 datasets for each degree were generated. **C.** Cross-comparison between DBN and CBN. 100 datasets were generated from the CBN model and 100 datasets were generated from the DBN model. Then the DBN and CBN models were fitted to all of the datasets, and the mean MCC in each class are reported.

Next, we compare the DP algorithm to CAPRI (Ramazzotti et al., 2015). CAPRI is a flexible framework for inferring cancer progression networks which can account for many types of interactions between nodes. CAPRI first applies a constraint-based algorithm to obtain a *prima facie* network, and then applies a local search algorithm to prune spurious edges. CAPRI is available through TRONCO De Sano et al. (2016). Figure 5A compares the ability of CAPRI and the DP algorithm to recover networks with various levels of noise. To understand how the algorithms perform as network complexity increases, Figure 5B varies the average degree while keeping the noise constant at *η* = 0.05.

Finally, we perform a cross-comparison of the CBN and DBN models. To do this, we simulate 100 datasets from the CBN model and 100 datasets from the DBN model. We fit the DBN model to the CBN datasets and vice verse. Figure 5C reports the mean MCC in each category.

### 3.2. Real Data Experiment

We use our method to recover the order of *driver* mutations in three cancer types from The Cancer Genome Atlas (TCGA) program (Cancer Genome Atlas Research Network et al., 2013). We selected Skin Cutaneous Melanoma (SKCM) and Lung Adenocarcinoma (LUAD) because there are known molecular subtypes and mutual exclusivity relationships characterized for them. To determine the driver mutations, we used results from (Bailey et al., 2018) where 26 computational methods had been applied to the TCGA data. The number of resulted driver mutations for SKCM and LUAD are below 30 and therefore exact DP method of Section 2.2 is applicable. We chose the Bladder Cancer (BLCA) for our third experiment since it has the highest driver mutation rate per sample in the TCGA data set (Bailey et al., 2018) and therefore is suitable to check the scalability of our proposed genetic algorithm. Number of samples, driver mutations, and frequent driver mutations (5% frequency cutoff threshold) for each cancer type is listed in Table 1.

**Table 1:** Number of samples (*n*), number of driver mutations, and number of frequent driver mutations (5% frequency cutoff threshold) (*p*) for the three used TCGA cancer types.

|  | # of samples | # of drivers | # of frequent drivers |
| --- | --- | --- | --- |
| **SKCM** | 467 | 20 | 15 |
| **LUAD** | 567 | 24 | 14 |
| **BLCA** | 414 | 45 | 31 |

To quantify our uncertainty in the estimated progression network, we run the algorithm on 100 bootstrapped datasets. We form the *mean graph* by only reporting the edges that are present in a sufficiently large number of networks estimated from the bootstrapped datasets (this cutoff will be 25 or 50).

#### 3.2.1. Progression of Mutations in Cutaneous Melanoma and Lung Adenocarcinoma

We run the DP method of the OncoBN package on 100 bootstrapped datasets with the in-degree bound of *k* = 3 and fixed universal spontaneous activation probability of *ε* = 0.025. The mean progression network is illustrated in Figure 6. Note that out of 24 LUAD mutations, only 11 of them are present in the mean progression network. This is because the rest of mutation are not connected with enough confident to the other nodes or to each others.

**Figure 6:**
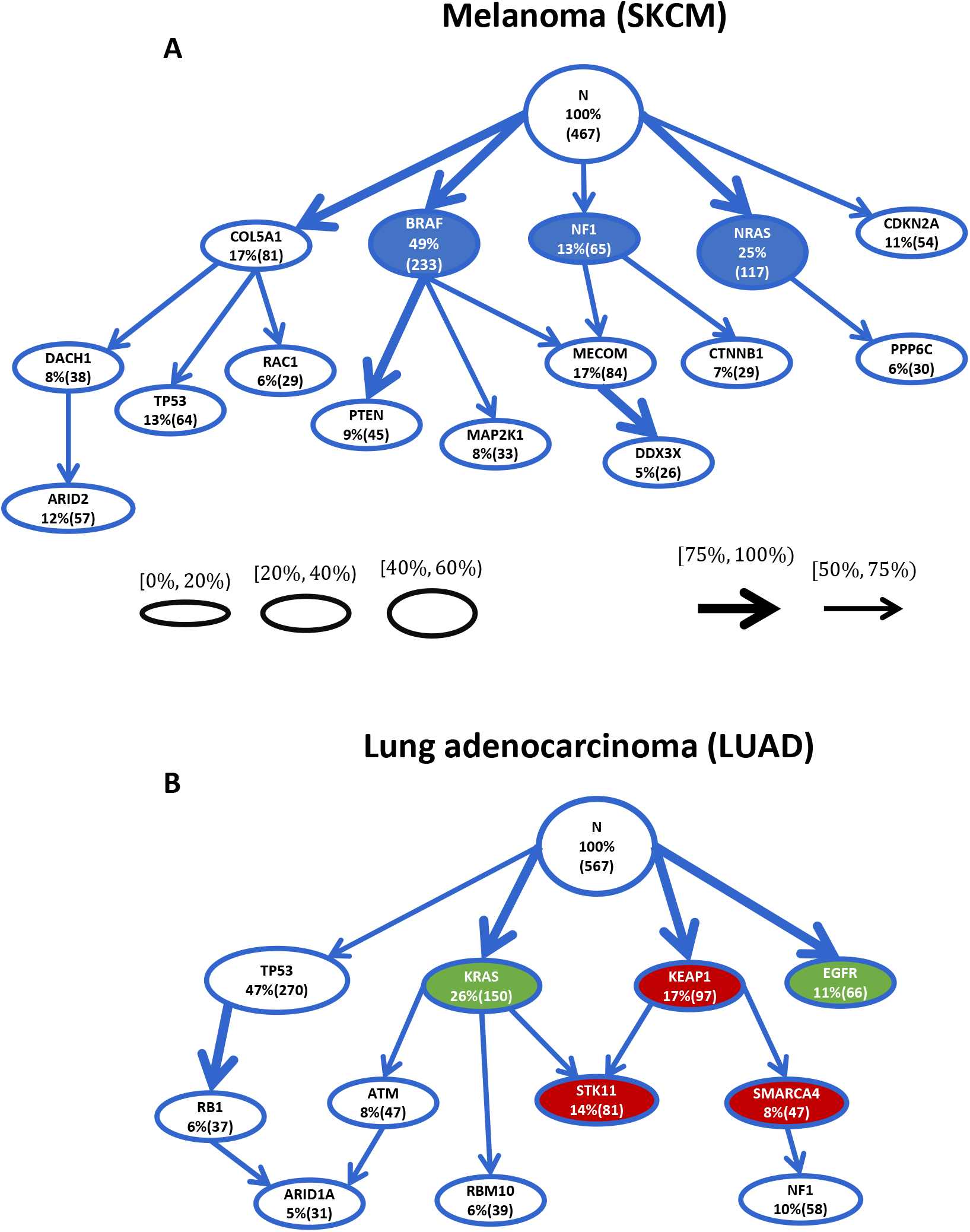
Mutation progression networks of melanoma and lung adenocarcinoma inferred by the exact dynamic programming learning method. **A.** We recover three known subtypes of melanoma (*BRAF*, *NRAS*, and *NF1* in blue) as separate roots. Mutations linked to metastasis such as *PTEN* and *DDX3X* are captured as late events. **B.** Synthetically lethal mutations of LUAD, *KRAS* and *EGFR* (green nodes), appear in disjoint branches. Frequently co-occurred mutations *STK11*, *KEAP1*, and *SMARCA4* occupy a branch of the inferred network (red nodes). Subtype defining mutations *TP53* and *RB1* are ordered with high confident.

For SKCM, we recovered three root mutations with a high mean presence: *BRAF*, *NRAS*, and *COL5A1*. In the rest of the graph, two connections have highest confident: *BRAF→PTEN* and *MECOM→DDX3X*. The only mutation with multiple parents is *MECOM*. For LUAD, three high confident roots have been recovered: *KRAS*, *KEAP1*, and *EGFR* plus a high confident edge *TP53→RB1*. *STK11* and *ARID1A* each have two parents.

#### 3.2.2. Progression of Mutations in Bladder Cancer

We run the GA of OncoBN package with 2*S* = 100 solutions for *T* = 300 generations on 100 bootstrap data sets. The mean progression network is illustrated in Figure 7. Out of *p* = 31 nodes, only 18 are inferred in the mean progression network because the remaining 13 are not connected with enough confident to the rest or to each others.

**Figure 7:**
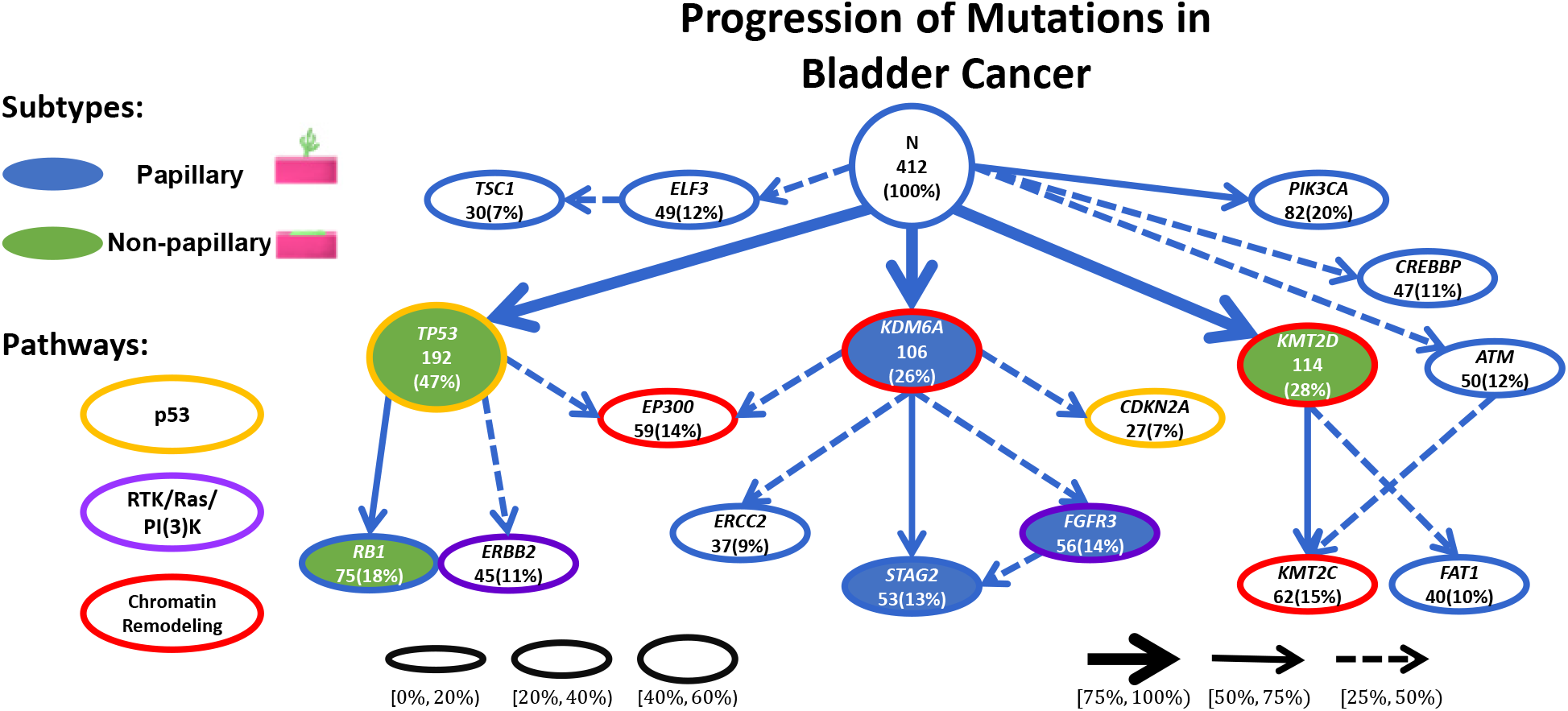
Mutation progression network of bladder cancer inferred by OncoBN. Focusing on the three high confidence roots (*TP53*, *KDM6A*, and *KMT2D*), the two subtypes of bladder cancer are clearly separated. The middle subgraph (rooted in *KDM6A*) is enriched for hallmark aberrations of the papillary subtype (blue nodes), and the other two subgraphs correspond to flat tumors (green nodes). Known mutual exclusive alteration pairs such as (*KDM6A*, *KMT2D*) and (*TP53*, *CDKN2A*) are occurring in different subgraphs. Four established highly perturbed pathways of bladder cancer are represented with varying outline colors. Each subtype has at least one mutated gene from these pathways is its subgraphs, therefore in both subtypes, all of the four pathways are perturbed.

We recover three root mutations with a high mean presence for the progression of bladder cancer: *TP53*, *KDM6A*, and *KMT2D*. From the several children of these roots, three have a mean presence greater than 50%: *RB1*, *STAG2*, and *KMT2C*. Finally, roots with meager mean presence (*ELF3*, *ATM*, and *CREBBP*) and childless *PIK3CA* are mutations for which OncoBN can not find enough supporting evidence to place them in the main progression graph. Note that these placements are possible because of the flexibility of the spontaneous activation model.

## 4. Discussion

### 4.1. Simulation Study

Figure 5 shows that the DP algorithm outperforms existing cancer progression algorithms when the noise rate is small. For high noise rates (*η ≈* 0.2), CAPRI is slightly more accurate. A future improvement to the method could be to integrate some of CAPRI’s regularization steps to improve robustness to noise. When *η* is small and fixed, DP uniformly outperforms CAPRI at different levels of network complexity (Figure 5B).

The cross-comparison (Figure 5C) shows that the DBN model is adequate even when the underlying data generating process assumes the CBN model. Although the CBN model performs slightly better when the data generating process assumes the CBN model (MCC 0.585 vs. MCC 0.44), the DBN model is significantly better when the data generating process assumes the DBN model (MCC 0.749 vs. MCC 0.448).

### 4.2. Melanoma and Lung Adenocarcinoma

Inferred melanoma progression network captures multiple known characteristics of melanoma. Namely, there are three distinct known molecular subtypes for cutaneous melanoma with *BRAF*, *NRAS*, and *NF1* as biomarkers (The Cancer Genome Atlas Network, 2015). All three of these mutations are roots of our inferred progression network, which suggests that they are important early occurring events. Strong metastasis inducing cooperation of *PTEN* with *BRAF* (Dankort et al., 2009) is captured with *BRAF→PTEN*. *DDX3X* that is linked with metastasis in melanoma is captured as a late stage event (Phung et al., 2019).

In the inferred progression network of lung adenocarcinoma, synthetically lethal mutations *KRAS* and *EGFR* (Unni et al., 2015) appear as distinct roots. Moreover, *KRAS*, *KEAP1*, *STK11*, *SMARCA4*, and *NF1* form a subgraph. It is known that *KRAS*, *KEAP1*, *STK11* and *SMARCA4* cooccur in non-small cell lung cancers (Schoenfeld et al., 2020) and our algorithm suggests that *KRAS* and *KEAP1* are early events in those tumors.

### 4.3. Bladder Cancer

The recovered progression network for bladder cancer reflects existing biological research. First, bladder cancer is known to have two histologically different subtypes known as papillary and non-papillary (Kamat et al., 2016). Papillary tumors are finger-like, which start in the lining and grow toward the center of the bladder. Non-papillary tumors also initiate in the lining but are flat in shape. Both types can be muscle-invasive, which means the tumor has grown outward, escaped the lining, and infiltrated bladder muscles, or non-muscle invasive (Kamat et al., 2016). All of the bladder cases in TCGA are muscle-invasive, but papillary and non-papillary cases are not known. There are known molecular signatures for papillary and non-papillary bladder cancers. Mutations in *TP53*, *RB1*, and *KMT2D* (green nodes in Figure 7) are very frequent in non-papillary sub-type while *KDM6A*, *STAG2*, and *FGFR3* (blue nodes in Figure 7) are hallmarks of papillary tumors (Dinney et al., 2004; Cancer Genome Atlas Research Network, 2014; Gui et al., 2011; Solomon et al., 2013). Focusing on the high confident recovered roots (*TP53*, *KDM6A*, and *KMT2D*) and their descendants, our inferred network of Figure 7 shows separate progression paths for papillary and non-papillary subtypes. The middle sub-graph rooted at *KDM6A* contains *KDM6A*, *STAG2*, and *FGFR3* mutations and is mostly separated from the rest of the network. Therefore we can match it to the progression of the papillary subtype. Sub-graphs on the right and left of the figure (rooted at *TP53* and *KMT2D*) are enriched with molecular hallmarks of non-papillary subtype. Our result shows the ability of OncoBN to infer the cancer progression network while maintaining subtype-specific biology.

In addition, we know that usually single perturbation of a pathway is enough for the manifestation of a cancer hallmark. Therefore, another mutated gene in the same pathway does not confer a selective advantage. Thus, patterns of mutual exclusivity of cancer events arise among genes in the same pathways. In bladder cancer, high rate of alteration of p53/Rb, RTK/Ras/PI(3)K, and histone modification pathways are observed (Cancer Genome Atlas Research Network, 2014). Figure 7 highlights the corresponding pathways of genes with different outline color for each pathway. It confirms that the two subtypes (papillary and non-papillary) both have perturbation in p53, RTK/Ras/PI(3)K, methylation, and acetylation pathways. The only mutation that is shared between the two subtypes is *EP300*, which corresponds to acetylation.

## References

Philipp M Altrock, Lin L Liu, and Franziska Michor. The mathematics of cancer: integrating quantitative models. Nat. Rev. Cancer, 15(12):730–745, December 2015.

Matthew H Bailey, Collin Tokheim, Eduard Porta-Pardo, Sohini Sengupta, Denis Bertrand, Amila Weerasinghe, Antonio Colaprico, Michael C Wendl, Jaegil Kim, Brendan Reardon, Patrick Kwok-Shing Ng, Kang Jin Jeong, Song Cao, Zixing Wang, Jianjiong Gao, Qingsong Gao, Fang Wang, Eric Minwei Liu, Loris Mularoni, Carlota Rubio-Perez, Niranjan Nagarajan, Isidro Cortés-Ciriano, Daniel Cui Zhou, Wen-Wei Liang, Julian M Hess, Venkata D Yellapantula, David Tamborero, Abel Gonzalez-Perez, Chayaporn Suphavilai, Jia Yu Ko, Ekta Khurana, Peter J Park, Eliezer M Van Allen, Han Liang, MC3 Working Group, Cancer Genome Atlas Research Network, Michael S Lawrence, Adam Godzik, Nuria Lopez-Bigas, Josh Stuart, David Wheeler, Gad Getz, Ken Chen, Alexander J Lazar, Gordon B Mills, Rachel Karchin, and Li Ding. Comprehensive characterization of cancer driver genes and mutations. Cell, 173(2):371–385.e18, April 2018.

David Barber. Bayesian reasoning and machine learning. Cambridge University Press, 2012.

Niko Beerenwinkel, Jörg Rahnenführer, Martin Däumer, Daniel Hoffmann, Rolf Kaiser, Joachim Selbig, and Thomas Lengauer. Learning multiple evolutionary pathways from cross-sectional data. J. Comput. Biol., 12(6):584–598, July 2005a.

Niko Beerenwinkel, Jörg Rahnenführer, Rolf Kaiser, Daniel Hoffmann, Joachim Selbig, and Thomas Lengauer. Mtreemix: a software package for learning and using mixture models of mutagenetic trees. Bioinformatics, 21(9):2106–2107, May 2005b.

Niko Beerenwinkel, Nicholas Eriksson, and Bernd Sturmfels. Conjunctive bayesian networks. Bernoulli, 13(4):893–909, November 2007.

Cancer Genome Atlas Research Network. Comprehensive molecular characterization of urothelial bladder carcinoma. Nature, 507(7492):315–322, March 2014.

Cancer Genome Atlas Research Network, John N Weinstein, Eric A Collisson, Gordon B Mills, Kenna R Mills Shaw, Brad A Ozenberger, Kyle Ellrott, Ilya Shmulevich, Chris Sander, and Joshua M Stuart. The cancer genome atlas Pan-Cancer analysis project. Nat. Genet., 45 (10):1113–1120, October 2013.

Giulio Caravagna, Alex Graudenzi, Daniele Ramazzotti, Rebeca Sanz-Pamplona, Luca De Sano, Giancarlo Mauri, Victor Moreno, Marco Antoniotti, and Bud Mishra. Algorithmic methods to infer the evolutionary trajectories in cancer progression. Proc. Natl. Acad. Sci., 113(28):E4025–34, July 2016.

Arthur Carvalho. A cooperative coevolutionary genetic algorithm for learning bayesian network structures. May 2013.

Yu-Kang Cheng, Rameen Beroukhim, Ross L Levine, Ingo K Mellinghoff, Eric C Holland, and Franziska Michor. A mathematical methodology for determining the temporal order of pathway alterations arising during gliomagenesis. PLoS Comput. Biol., 8(1):e1002337, January 2012.

Simona Cristea, Jack Kuipers, and Niko Beerenwinkel. pathTiMEx: Joint inference of mutually exclusive cancer pathways and their progression dynamics. J. Comput. Biol., 24(6):603–615, June 2017.

Ibiayi Dagogo-Jack and Alice T. Shaw. Tumour heterogeneity and resistance to cancer therapies. Nature Reviews Clinical Oncology, 15(2):81–94, 2018.

David Dankort, David P. Curley, Robert A. Cartlidge, Betsy Nelson, Anthony N. Karnezis, William E. Damsky, Mingjian J. You, Ronald A. DePinho, Martin McMahon, and Marcus Bosenberg. Braf(V600E) cooperates with Pten loss to induce metastatic melanoma. Nature Genetics, 41(5):544–552, 2009.

Alexander Davis, Ruli Gao, and Nicholas Navin. Tumor evolution: Linear, branching, neutral or punctuated? Biochimica Et Biophysica Acta. Reviews on Cancer, 1867(2):151–161, 2017.

Luca De Sano, Giulio Caravagna, Daniele Ramazzotti, Alex Graudenzi, Giancarlo Mauri, Bud Mishra, and Marco Antoniotti. TRONCO: an R package for the inference of cancer progression models from heterogeneous genomic data. Bioinformatics, 32(12):1911–1913, June 2016.

Yulan Deng, Shangyi Luo, Chunyu Deng, Tao Luo, Wenkang Yin, Hongyi Zhang, Yong Zhang, Xinxin Zhang, Yujia Lan, Yanyan Ping, Yun Xiao, and Xia Li. Identifying mutual exclusivity across cancer genomes: Computational approaches to discover genetic interaction and reveal tumor vulnerability. Briefings in Bioinformatics, 20(1):254–266, 2019.

R Desper, F Jiang, O P Kallioniemi, H Moch, C H Papadimitriou, and A A Schäffer. Inferring tree models for oncogenesis from comparative genome hybridization data. J. Comput. Biol., 6(1): 37–51, 1999.

Colin P N Dinney, David J McConkey, Randall E Millikan, Xifeng Wu, Menashe Bar-Eli, Liana Adam, Ashish M Kamat, Arlene O Siefker-Radtke, Tomasz Tuziak, Anita L Sabichi, H Barton Grossman, William F Benedict, and Bogdan Czerniak. Focus on bladder cancer. Cancer Cell, 6(2):111–116, August 2004.

Hossein Shahrabi Farahani and Jens Lagergren. Learning Oncogenetic Networks by Reducing to Mixed Integer Linear Programming. PLOS ONE, 8(6):e65773, 2013.

E R Fearon and B Vogelstein. A genetic model for colorectal tumorigenesis. Cell, 61(5):759–767, June 1990.

Moritz Gerstung, Michael Baudis, Holger Moch, and Niko Beerenwinkel. Quantifying cancer progression with conjunctive bayesian networks. Bioinformatics, 25(21):2809–2815, November 2009.

Moritz Gerstung, Nicholas Eriksson, Jimmy Lin, Bert Vogelstein, and Niko Beerenwinkel. The temporal order of genetic and pathway alterations in tumorigenesis. PLoS One, 6(11):e27136, November 2011.

Moritz Gerstung, Clemency Jolly, Ignaty Leshchiner, Stefan C. Dentro, Santiago Gonzalez, Daniel Rosebrock, Thomas J. Mitchell, Yulia Rubanova, Pavana Anur, Kaixian Yu, Maxime Tarabichi, Amit Deshwar, Jeff Wintersinger, Kortine Kleinheinz, Ignacio Vázquez-García, Kerstin Haase, Lara Jerman, Subhajit Sengupta, Geoff Macintyre, Salem Malikic, Nilgun Donmez, Dimitri G. Livitz, Marek Cmero, Jonas Demeulemeester, Steven Schumacher, Yu Fan, Xiaotong Yao, Juhee Lee, Matthias Schlesner, Paul C. Boutros, David D. Bowtell, Hongtu Zhu, Gad Getz, Marcin Imielinski, Rameen Beroukhim, S. Cenk Sahinalp, Yuan Ji, Martin Peifer, Florian Markowetz, Ville Mustonen, Ke Yuan, Wenyi Wang, Quaid D. Morris, Paul T. Spellman, David C. Wedge, and Peter Van Loo. The evolutionary history of 2,658 cancers. Nature, 578(7793): 122–128, 2020.

Yaoting Gui, Guangwu Guo, Yi Huang, Xueda Hu, Aifa Tang, Shengjie Gao, Renhua Wu, Chao Chen, Xianxin Li, Liang Zhou, Minghui He, Zesong Li, Xiaojuan Sun, Wenlong Jia, Jinnong Chen, Shangming Yang, Fangjian Zhou, Xiaokun Zhao, Shengqing Wan, Rui Ye, Chaozhao Liang, Zhisheng Liu, Peide Huang, Chunxiao Liu, Hui Jiang, Yong Wang, Hancheng Zheng, Liang Sun, Xingwang Liu, Zhimao Jiang, Dafei Feng, Jing Chen, Song Wu, Jing Zou, Zhongfu Zhang, Ruilin Yang, Jun Zhao, Congjie Xu, Weihua Yin, Zhichen Guan, Jiongxian Ye, Hong Zhang, Jingxiang Li, Karsten Kristiansen, Michael L Nickerson, Dan Theodorescu, Yingrui Li, Xiuqing Zhang, Songgang Li, Jian Wang, Huanming Yang, Jun Wang, and Zhiming Cai. Frequent mutations of chromatin remodeling genes in transitional cell carcinoma of the bladder. Nat. Genet., 43(9):875–878, August 2011.

Douglas Hanahan and Robert A. Weinberg. Hallmarks of Cancer: The Next Generation. Cell, 144 (5):646–674, 2011. ISSN 0092-8674, 1097-4172. doi: 10.1016/j.cell.2011.02.013.

Markus Kalisch, Martin Mächler, Diego Colombo, Marloes H Maathuis, and Peter Bühlmann. Causal inference using graphical models with the r package pcalg. Journal of Statistical Software, 47(11):1–26, 2012.

Ashish M Kamat, Noah M Hahn, Jason A Efstathiou, Seth P Lerner, Per-Uno Malmström, Woonyoung Choi, Charles C Guo, Yair Lotan, and Wassim Kassouf. Bladder cancer. The Lancet, 388 (10061):2796–2810, December 2016.

Daphne Koller and Nir Friedman. Probabilistic graphical models: principles and techniques. MIT press, 2009.

Wai Lam and Fahiem Bacchus. LEARNING BAYESIAN BELIEF NETWORKS: AN APPROACH BASED ON THE MDL PRINCIPLE. Comput. Intell., 10(3):269–293, August 1994.

Mark D M Leiserson, Hsin-Ta Wu, Fabio Vandin, and Benjamin J Raphael. CoMEt: a statistical approach to identify combinations of mutually exclusive alterations in cancer. Genome Biol., 16: 160, August 2015.

Brian W Matthews. Comparison of the predicted and observed secondary structure of t4 phage lysozyme. Biochimica et Biophysica Acta (BBA)-Protein Structure, 405(2):442–451, 1975.

Hesam Montazeri, Jack Kuipers, Roger Kouyos, Jürg Böni, Sabine Yerly, Thomas Klimkait, Vincent Aubert, Huldrych F Günthard, Niko Beerenwinkel, and Swiss HIV Cohort Study. Large-scale inference of conjunctive bayesian networks. Bioinformatics, 32(17):i727–i735, 2016a.

Hesam Montazeri, Jack Kuipers, Roger Kouyos, Jürg Böni, Sabine Yerly, Thomas Klimkait, Vincent Aubert, Huldrych F Günthard, Niko Beerenwinkel, and Swiss HIV Cohort Study. Large-scale inference of conjunctive bayesian networks. Bioinformatics, 32(17):i727–i735, September 2016b.

Loes Olde Loohuis, Giulio Caravagna, Alex Graudenzi, Daniele Ramazzotti, Giancarlo Mauri, Marco Antoniotti, and Bud Mishra. Inferring tree causal models of cancer progression with probability raising. PLoS One, 9(10):e108358, October 2014.

Bengt Phung, Maciej Cieśla, Adriana Sanna, Nicola Guzzi, Giulia Beneventi, Phuong Cao Thi Ngoc, Martin Lauss, Rita Cabrita, Eugenia Cordero, Ana Bosch, Frida Rosengren, Jari Häkkinen, Klaus Griewank, Annette Paschen, Katja Harbst, Haakan Olsson, Christian Ingvar, Ana Carneiro, Hensin Tsao, Dirk Schadendorf, Kristian Pietras, Cristian Bellodi, and Göran Jönsson. The X-Linked DDX3X RNA Helicase Dictates Translation Reprogramming and Metastasis in Melanoma. Cell Reports, 27(12):3573–3586.e7, 2019.

Daniele Ramazzotti, Giulio Caravagna, Loes Olde Loohuis, Alex Graudenzi, Ilya Korsunsky, Giancarlo Mauri, Marco Antoniotti, and Bud Mishra. CAPRI: efficient inference of cancer progression models from cross-sectional data. Bioinformatics, 31(18):3016–3026, September 2015.

Daniele Ramazzotti, Alex Graudenzi, Giulio Caravagna, and Marco Antoniotti. Modeling cumulative biological phenomena with Suppes-Bayes causal networks. Evol. Bioinform. Online, 14: 1176934318785167, July 2018.

Benjamin J Raphael and Fabio Vandin. Simultaneous inference of cancer pathways and tumor progression from cross-sectional mutation data. J. Comput. Biol., 22(6):510–527, June 2015.

Adam J. Schoenfeld, Chai Bandlamudi, Jessica A. Lavery, Joseph Montecalvo, Azadeh Namakydoust, Hira Rizvi, Jacklynn Egger, Carla P. Concepcion, Sonal Paul, Maria E. Arcila, Yahya Daneshbod, Jason Chang, Jennifer L. Sauter, Amanda Beras, Marc Ladanyi, Tyler Jacks, Charles M. Rudin, Barry S. Taylor, Mark T. A. Donoghue, Glenn Heller, Matthew D. Hellmann, Natasha Rekhtman, and Gregory J. Riely. The Genomic Landscape of SMARCA4 Alterations and Associations with Outcomes in Patients with Lung Cancer. Clinical Cancer Research, 26 (21):5701–5708, 2020.

Tomi Silander and Petri Myllymäki. A simple approach for finding the globally optimal Bayesian network structure. In Proceedings of the Twenty-Second Conference on Uncertainty in Artificial Intelligence, pages 445–452, 2006.

David A Solomon, Jung-Sik Kim, Jolanta Bondaruk, Shahrokh F Shariat, Zeng-Feng Wang, Abdel G Elkahloun, Tomoko Ozawa, Julia Gerard, Dazhong Zhuang, Shizhen Zhang, Neema Navai, Arlene Siefker-Radtke, Joanna J Phillips, Brian D Robinson, Mark A Rubin, Björn Volkmer, Richard Hautmann, Rainer Küfer, Pancras C W Hogendoorn, George Netto, Dan Theodorescu, C David James, Bogdan Czerniak, Markku Miettinen, and Todd Waldman. Frequent truncating mutations of STAG2 in bladder cancer. Nat. Genet., 45(12):1428–1430, December 2013.

The Cancer Genome Atlas Network. Genomic Classification of Cutaneous Melanoma. Cell, 161 (7):1681–1696, 2015.

Arun M Unni, William W Lockwood, Kreshnik Zejnullahu, Shih-Queen Lee-Lin, and Harold Varmus. Evidence that synthetic lethality underlies the mutual exclusivity of oncogenic KRAS and EGFR mutations in lung adenocarcinoma. eLife, 4:e06907, 2015.

