## Supplement for "Oncogenetic Network Estimation with Disjunctive Bayesian Networks"

---

---

#### A. Measurement Error Model

One can attribute deviation from the progression network to measurement error, i.e., presence of false positive and negative observations (Szabo and Boucher, 2002). False positives and negatives can arise from errors in measurement technology. A false negative can also arise from having a single sample from a spatially heterogeneous tumor. We assume that there are unique false positive $\xi^+$ and negative $\xi^-$ probabilities that generates the observed event $\mathbf{x}$ from the underlying latent event $\mathbf{z}$ as: $\mathbb{P}(x_j = 1|z_j = 0) = \xi^+, \mathbb{P}(x_j = 0|z_j = 1) = \xi^-$, Figure 1.

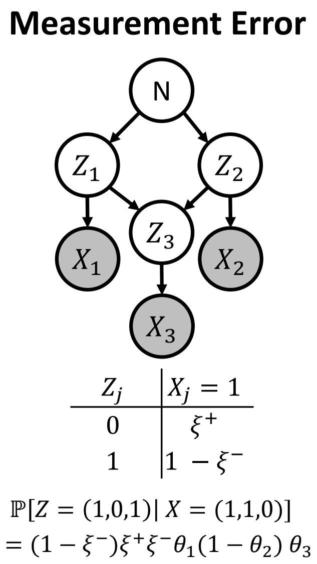


Figure 1: **Bayesian networks of the three cancer progression models investigated.** Node $N$ represents normal cell state, and each random variable $X_j$ is an observed alteration, and the corresponding progression probability parameter is $\theta_j$. The conditional probability table of $X_3$ is shown, and probabilities of an instance observation is computed. Measurement error model where each actual unobserved alteration $Z_j$ generates an observation $X_j$ according to universal false positive and negative probabilities.

### B. Parameter Estimation

Given $n$ cross-sectional samples and the progression network $G$, we wish to find $\hat{\boldsymbol{\theta}}_G^{\mathrm{MLE}}$, the maximum likelihood estimator (MLE) for $\boldsymbol{\theta}$ in each of the three variants of the DBN. The likelihood and MLE for the spontaneous activation model was presented in the paper, here, we discuss all MLE for all three models.

#### *B.1. MLE for the Basic DBN*

First, we write the joint distribution of events using matrix $\mathbf{A}$ (the adjacency matrix of $G$) compactly.

**Proposition 1.** *The likelihood of the basic DBN can be written as*

$$\mathbb{P}(\mathbf{x};\boldsymbol{\theta},G)=\prod_{j=1}^{p}[\theta_j{}^{x_j}(1-\theta_j)^{1-x_j}]^{\mathbf{1}(\mathbf{x}(\mathcal{P}_\mathbf{j})\neq\mathbf{0})}(1-x_j)^{\mathbf{1}(\mathbf{x}(\mathcal{P}_\mathbf{j})=\mathbf{0})} \tag{1}$$

*where* $\mathbf{1}$ *is the indicator function and* $\mathbf{1}(\mathbf{x}(\mathcal{P}_\mathbf{j})\neq\mathbf{0})$ *checks if any of* $j$*'s parents has occurred. Note that* $\mathbf{x}(\mathcal{P}_j)$ *can be computed easily as* $(\mathbf{a}_j\odot\mathbf{x})_{\mathcal{P}_j}$ *where* $\mathbf{a}_j$ *is the* $j$*th column of* $\mathbf{A}$*,* $\odot$ *is the Hadamard product and the subscript* $\mathcal{P}_j$ *selects parents of* $j$ *from the vector.*

Using the compact representation (1) of the likelihood, we can compute the MLE of $\boldsymbol{\theta}$.

**Proposition 2.** *Given* $n$ *independent samples* $\{\mathbf{x}_i\}_{i=1}^n$ *from the same population defined by* $G$ *and* $\theta$ *where* $\mathbf{x}_i\in\{0,1\}^p$*, MLE for* $\theta_j$ *is*

$$\hat{\theta}_j^{MLE}=\frac{\sum_{i=1}^n\mathbf{1}(\mathbf{x_{ij}}=\mathbf{1},\mathbf{x_i}(\mathcal{P}_\mathbf{j})\neq\mathbf{0})}{\sum_{i=1}^n\mathbf{1}(\mathbf{x_i}(\mathcal{P}_\mathbf{j})\neq\mathbf{0})}. \tag{2}$$

*where* $x_{ij}$ *is the realization of the* $j$*th event in the* $i$*th sample.*

Intuitively, (2) is just a sample proportion. The denominator counts the number of samples in which at least one of the parents of $j$ occurred while the numerator counts those where $j$ occurred along with at least one of its parents.

#### *B.2. MLE for the Spontaneous Activation Model.*

The likelihood of the spontaneous activation model is as follows:

$$\mathbb{P}(\mathbf{x};\boldsymbol{\theta},G)=\prod_{j=1}^{p}[\theta_j{}^{x_j}(1-\theta_j)^{1-x_j}]^{\mathbf{1}\left(\mathbf{x}(\mathcal{P}_\mathbf{j})\neq\mathbf{0}\right)}\varepsilon_j^{\mathbf{1}\left(\mathbf{x}(\mathcal{P}_\mathbf{j})=\mathbf{0}\right)}. \tag{3}$$

Similarities between likelihoods (1) and (3) suggest that the MLE for $\boldsymbol{\theta}$ of the spontaneous activation model should be the same as for the $\boldsymbol{\theta}$ of the basic DBN presented in (2). So, we only compute the MLE for $\varepsilon$.

**Proposition 3.** *Given $n$ independent samples from the same population defined by $G$ and $\theta$, the MLE of $\theta$ for the spontaneous activation model is as (2) and the MLE of $\varepsilon_j$ can be computed as:*

$$\hat{\varepsilon}_j^{MLE} = \frac{\sum_{i=1}^{n} \mathbf{1}(\mathbf{x_{ij}} = \mathbf{1}, \mathbf{x_i}(\mathcal{P}_\mathbf{j}) = \mathbf{0})}{\sum_{i=1}^{n} \mathbf{1}(\mathbf{x_i}(\mathcal{P}_\mathbf{j}) = \mathbf{0})}. \tag{4}$$

*where $x_{ij}$ is the realization of the $j$th event in the $i$th sample.*

The ratio in (4) counts the percentage of samples in which $j$ has occurred without any parent and is thus an intuitively reasonable estimator of the spontaneous activation rate.

#### *B.3. EM for the Measurement Error Model*

Assuming $\xi^+$ and $\xi^-$ are fixed and known, the MLE of $\boldsymbol{\theta}$ can be approximated using the Expectation Maximization (EM) algorithm. Given the $t$th EM iteration estimate $\boldsymbol{\theta}^{(t)}$ for $\boldsymbol{\theta}$, we set

$$\boldsymbol{\theta}^{(t+1)} = \underset{\boldsymbol{\theta}}{\operatorname{argmax}} \sum_{i=1}^{n} \sum_{\mathbf{z}_i} \mathbb{P}(\mathbf{z}_i|\mathbf{x}_i; \boldsymbol{\theta}^{(t)}, \xi^+, \xi^-)\ell(\boldsymbol{\theta}; \mathbf{x}_i, \mathbf{z}_i), \tag{5}$$

where $\ell(\boldsymbol{\theta}; \mathbf{x}_i, \mathbf{z}_i) = \log \mathbb{P}(\mathbf{x}_i, \mathbf{z}_i; \boldsymbol{\theta})$ is the joint log-likelihood of sample $i$. The update for $\boldsymbol{\theta}^{(t)}$ can be found explicitly as follows.

**Theorem 1** (Closed form EM Update)**.** *Given $n$ independent samples $\{\mathbf{x}_i\}_{i=1}^n$ from the same population defined by $G$ and $\theta$ where $\mathbf{x}_i \in \{0,1\}^p$, the EM update (7) for $\theta_j^{(t+1)}$ has the following closed form:*

$$\theta_j^{(t+1)} = \frac{\sum_{i=1}^{n} \sum_{\mathbf{z}_i} \mathbf{1}(\mathbf{z_{ij}} = \mathbf{1}, \mathbf{z_i}(\mathcal{P}_\mathbf{j}) \neq \mathbf{0})\mathbb{P}(\mathbf{z_i}|\mathbf{x_i})}{\sum_{i=1}^{n} \sum_{\mathbf{z}_i} \mathbf{1}(\mathbf{z_i}(\mathcal{P}_\mathbf{j}) \neq \mathbf{0})\mathbb{P}(\mathbf{z_i}|\mathbf{x_i})}. \tag{6}$$

Note that if there is no measurement error, i.e., $\xi^+ = \xi^- = 0$, then (6) reduces to (2). In practice, computing (6) incurs exponential time because the inner sum goes through all $2^p$ possible realizations of the latent vector $\mathbf{z}_i$. Consequently, this model is only practically useful when $p \leq 12$.

#### *B.4. Proofs*

##### *B.4.1. Proofs of Propositions 2 and 3*

The propositions are special cases of maximum likelihood estimation of Table-CPDs for a given Bayesian Network (BN) presented in Section 17.2.3 of Koller and Friedman (2009).

##### *B.4.2. Proof of Theorem 1*

*Proof.* We should set the gradient of the following objective function to zero and solve for $\boldsymbol{\theta}$:

$$f(\boldsymbol{\theta}) = \sum_{i=1}^{n} \sum_{\mathbf{z}_i} \mathbb{P}(\mathbf{z}_i|\mathbf{x}_i; \boldsymbol{\theta}^{(t)}, \xi^+, \xi^-)\ell(\boldsymbol{\theta}; \mathbf{x}_i, \mathbf{z}_i), \tag{7}$$

The general form of this tedious calculation can be found in Chapter 19 of Koller and Friedman (2009). Instantiating the calculation in Section 19.2.2.3 of Koller and Friedman (2009) performed for general Table-CPDs for the specific family of DBN's CPDs completes the proof. ∎

#### C. Genetic Algorithm Operations for Structure Learning

We represent a DAG $G$ with a pair $(\mathbf{O}, \boldsymbol{\pi})$, where $\mathbf{O}$ is the adjacency matrix for the *topological ordering* of $G$ (i.e., an strictly upper triangular matrix), and $\boldsymbol{\pi}$ is a permutation vector for labeling the vertices. During each generation of the GA, we evolve $\mathbf{O}$ and $\boldsymbol{\pi}$ of each $G$ separately. Therefore, we need to define crossover and mutation operation for both ordering matrices and permutation vectors.

**Crossover.** Each crossover operation is defined to take in two DAGs and produce two offspring so that the number of individuals per generation remains constant. For the two selected DAGs their orderings and permutations are crossed over as follows.

- *Ordering Crossover*: With probability $c_o$, the two upper triangular matrix chromosomes are recombined by interchanging their rows.
- *Permutation Crossover*: With probability $c_\pi$, the permutation chromosomes are recombined using the cycle crossover algorithm Oliver et al. (1987).

If no crossover occurs, the two selected chromosomes are passed down to the next generation unchanged. **Mutation.** To maintain diversity in the population, we also define several mutation operators.

- *Edge Mutation*: With probability $m_e$, either an edge is added or an existing edge is removed which is achieved by randomly flipping elements on the upper triangle of $\mathbf{O}$.
- *Branch Mutation*: A branch is defined as a vertex along with all of its descendant vertices. With probability $m_b$, a branch is randomly selected and is moved to another location. This operation is done by cutting offs all parents of the root and assigning a random parent to it. Also, all parents of the other nodes in the branch that are not member of the branch themselves are cut off.
- *Permutation Mutation*: With probability $m_\pi$, two elements in the permutation chromosome $\boldsymbol{\pi}$ are swapped.

Edge and branch mutations operate on $\mathbf{O}$ and permutation mutation operates on $\boldsymbol{\pi}$.

### D. An Elaboration on DAG Equivalence Classes

Many different network structures induce the same probability distribution over $\mathbf{x} \in \{0,1\}^p$, i.e., likelihood.

**Definition 2.** We say that $G$ and $G'$ are equivalent and show it by $G \sim G'$ if for every $\boldsymbol{\theta}$ and $\mathbf{x}$, $\mathbb{P}(\mathbf{x}; G, \boldsymbol{\theta}) = \mathbb{P}(\mathbf{x}; G', \boldsymbol{\theta})$.

It is clear that $\sim$ defines an equivalence relation over DAGs. Since DAGs belonging to a same equivalence class are indistinguishable likelihood-wise (i.e., with current data) we can select a representative for each class and just search the space of all representatives to speed up the algorithm. We call this representative DAG the **canonical form** of the equivalence class. In the following, we will show how we can make the canonical form from the given graph $G$.

Remember that each DAG can be represented by an ordering $\mathbf{O}$ (upper triangular matrix) and a permutation vector $\boldsymbol{\pi}$, i.e., $G = (\mathbf{O}, \boldsymbol{\pi})$. To make the canonical form of graph $G$, one should remove *redundant edges* from $\mathbf{O}$ and give unique labels to *similar vertices* by $\boldsymbol{\pi}$. In the following, we show how to detect redundant edges and similar vertices and finally we prove that if $G$ and $G'$ have the same canonical form, they belong to the same equivalence class, i.e., represent the same likelihood.

**Definition 3.** For each event (node) $X_i$, define $\mathcal{C}_i$ and $\mathcal{P}_i$ as the set of children and parents of $i$ respectively.

#### *D.1. Redundant Edges*

Below, we define redundant edges as those that if removed they do not change the corresponding probability distribution of the BN.

**Definition 4** (**Redundant Edges**). An edge $e$ in a Bayesian Network structure $G$ is said to be **redundant** if the resulting BN structure $G'$ that is obtained by removing $e$ is equivalent to $G$, i.e., $G \sim G'$.

Next, we show how we can identify redundant edges of a given graph for various version of DBN models.

##### *D.1.1. Detecting Redundant Edges in Basic DBN and Measurement Error Models*

**Proposition 4.** *Let $e = (j, i)$ be an edge from node $j$ to node $i$ in DAG $G'$. Under the Basic DBN and the measurement error models, edge $e$ is redundant if and only if every path from the root node $N$ to $j$ passes through another parent of node $i$. Mathematically, let $\Phi$ be the set of all paths from the root $N$ to $j$. Then, $e$ is redundant if and only if $\forall \phi \in \Phi : \phi \cap \mathcal{P}_i \neq \emptyset$ where $\mathcal{P}_i$ is the set of parents of $i$ in $G$.*

| | $x_j$ | $\mathbf{x}(\mathcal{P}_i)$ | $\mathbb{P}(x_i = 1 \vert \mathbf{x}(\mathcal{P}_i), x_j)$ | $\mathbb{P}(x_i = 1 \vert \mathbf{x}(\mathcal{P}_i))$ |
|---|---|---|---|---|
| 1 | 0 | $=\mathbf{0}$ | 1 | 1 |
| 2 | 0 | $\neq \mathbf{0}$ | $\theta_i$ | $\theta_i$ |
| 3 | 1 | $=\mathbf{0}$ | $\mathbb{P}(x_i = 1 \vert \mathbf{x}(\mathcal{P}_i) = \mathbf{0}, x_j = 1)$ | 0 |
| 4 | 1 | $\neq \mathbf{0}$ | $\theta_i$ | $\theta_i$ |

Table 1: Comparing the local CPDs of node $i$ in $G'$ and $G$ where their only difference is the presence of the edge $e = (j, i)$ in $G'$.

*Proof.* The proof is similar for both models, so we use the notation of the Basic DBN model. Note that we want to compare the likelihood of graphs $G'$ and $G$ for the fixed sample $\mathbf{x}$ and fixed parameter $\theta$, where $G'$ has one extra edge $e$. For convenience we consider $\mathcal{P}_i$ as the parents of $i$ in $G$ and $\mathcal{P}'_i = \mathcal{P}_i \cup \{j\}$ as $i$'s parents in $G'$. The only difference between $\mathbb{P}(\mathbf{x}; G, \theta)$ and $\mathbb{P}(\mathbf{x}; G', \theta)$ is in the contribution of $j$ in the likelihood, i.e., $\mathbb{P}(x_i|\mathbf{x}_{\mathcal{P}_i})$ in $G$ vs. $\mathbb{P}(x_i|\mathbf{x}_{\mathcal{P}_i}, x_j)$ in $G'$. Table 1 shows the local CPDs of $i$ in $G$ and $G'$.

The only difference of the two conditional distribution is in the line three of the table. Distributions are equal if and only if $\mathbb{P}(x_i = 1|\mathbf{x}(\mathcal{P}_i) = \mathbf{0}, x_j = 1) = 0$. In other words, we need to make the $\{\mathbf{x}(\mathcal{P}_i) = \mathbf{0}, x_j = 1\}$ event impossible. To this end, whenever node $j$ is active ($x_j = 1$), at least one of the parents of $i$ must be active ($\mathbf{x}_{\mathcal{P}_i} \neq \mathbf{0}$).

We also know that in the Basic DBN model a node $j$ is active if and only if there is at least one path $\phi$ of active nodes from the root $N$ to $j$. Therefore, if there exists at least one parent of $i$ in every path $\phi$ from root to $j$ the $\{\mathbf{x}(\mathcal{P}_i) = \mathbf{0}, x_j = 1\}$ event will never occur which proves the proposition. ■

*D.1.2. Detecting Redundant Edges in the Spontaneous Activation Model*

Redundant edges in the spontaneous activation model are different from those of the basic DBN and the measurement error models. Below proposition characterizes redundant edges for the spontaneous activation model.

**Proposition 5.** *Let $e = (j, i)$ be an edge from node $j$ to node $i$ in DAG $G'$. Under the spontaneous activation model, edge $e$ is redundant if and only if every node in every path from the root node $N$ to $j$ is a parent of node $i$. Mathematically, let $\Phi$ be the set of all paths from the root $N$ to $j$. Then, $e$ is redundant if and only if $\forall \phi \in \Phi$ and $\forall k \in \phi : x_k \in \mathcal{P}_i$ where $\mathcal{P}_i$ is the set of parents of $i$ in $G$.*

*Proof.* The proof is similar to that of the Proposition 4 except in the last step where we want to make $\{\mathbf{x}(\mathcal{P}_i) = \mathbf{0}, x_j = 1\}$ event impossible. In the spontaneous activation model, node $j$ can become activate in three general ways: 1) by an active path from $N$ to $j$ or 2) by an active path from a node $k$ who is spontaneously activated, or 3) by self activation with probability $\varepsilon_j$. In the former case, the proof of Proposition 4 goes through. When activation of $j$ is due to spontaneous activation of itself or any of its ancestors, $\{\mathbf{x}(\mathcal{P}_i) = \mathbf{0}\}$ is possible unless all of the nodes in all of the paths from $N$ to $j$ be parents of $i$. ■

**Remark.** Note that due to the stringent definition of redundant edges for the spontaneous activation model (Proposition 5) there are not many of them in a given graph and therefore sizes of equivalence classes for the spontaneous activation model are small. As a result, the corresponding search space of canonical forms is large andthe speed gain of searching only canonical forms becomes scant. To avoid this issue, we use the definition of redundant edges of other models (basic and measurement error) for the spontaneous activation model. This approximation is justifiable since the spontaneous activation probability of each node is usually very small ($\varepsilon_j \leq 0.05$) and therefore any path resulting from spontaneous activation has a very low probability which makes $\{\mathbf{x}(\mathcal{P}_i) = \mathbf{0}, x_j = 1\}$ event *close* to impossible. With this approximation for spontaneous activation model we gain scalability without completely compromising the theoretical properties of the proposed algorithm.

*D.2. Similar Vertices*

Below, we define similar vertices as those that if their labels are swapped the corresponding probability distribution of the BN will not change.

**Definition 5** (**Similar Vertices**)**.** Nodes $i$ and $j$ of a Bayesian Network structure $G = (\mathbf{O}, \boldsymbol{\pi})$ are **similar** if the resulting BN structure $G' = (\mathbf{O}, \boldsymbol{\pi}')$ that is obtained by swapping $\pi_i$ and $\pi_j$ is equivalent to $G$, i.e., $G \sim G'$.

Next, we show how we can identify similar vertices of a given graph for all DBN models.

**Proposition 6.** *Let $G = (\mathbf{O}, \boldsymbol{\pi})$ and $G' = (\mathbf{O}, \boldsymbol{\pi}')$ be two DAGs with the same topological ordering $\mathbf{O}$. Let the set of parents and children of node $i$ in $G$ represented by $\mathcal{P}_i$ and $\mathcal{C}_i$ and in $G'$ by $\mathcal{P}'_i$ and $\mathcal{C}'_i$ respectively. Then $G \sim G'$ if and only if $\mathcal{P}_i = \mathcal{P}_j$ and $\mathcal{C}_i = \mathcal{C}_j$.*

*Proof.* Intuitively, note that local CPDs of nodes $i$, $j$, and their children stays the same if labels of $i$ and $j$ are swapped. Mathematically, the only contribution of $i$, $j$, and their children in the joint distribution of BN represented by $G$ are the following conditional probabilities:

1. $\mathbb{P}(x_{\pi_i} | \mathbf{x}(\mathcal{P}_i))$
2. $\mathbb{P}(x_{\pi_j} | \mathbf{x}(\mathcal{P}_j))$
3. $\forall k \in \mathcal{C}_i \cup \mathcal{C}_j : \mathbb{P}(x_{\pi_k} | \mathbf{x}(\mathcal{P}_k))$

Note that $k$ is the child of of $i$, $j$, or both of them.

To have the same probability distribution the product of these three class of CPDs should be equal to the product of their counterparts in $G'$, i.e., the following conditions must be satisfied:

$$\mathbb{P}(x_{\pi_i} | \mathbf{x}(\mathcal{P}_i)) \mathbb{P}(x_{\pi_j} | \mathbf{x}(\mathcal{P}_j)) \prod_{k \in \mathcal{C}_i \cup \mathcal{C}_j} \mathbb{P}(x_{\pi_k} | \mathbf{x}(\mathcal{P}_k)) = \mathbb{P}(x_{\pi'_i} | \mathbf{x}(\mathcal{P}'_i)) \mathbb{P}(x_{\pi'_j} | \mathbf{x}(\mathcal{P}'_j)) \prod_{k \in \mathcal{C}'_i \cup \mathcal{C}'_j} \mathbb{P}(x_{\pi_k} | \mathbf{x}(\mathcal{P}'_k))$$

Because of the swapping of $i$ and $j$'s labels, we have $\pi'_i = \pi_j$, $\pi'_j = \pi_i$ while $\mathcal{P}'_i = \mathcal{P}_i$, $\mathcal{P}'_j = \mathcal{P}_j$, $\mathcal{C}'_i = \mathcal{C}_i$, and $\mathcal{C}'_j = \mathcal{C}_j$ because $\mathbf{O}$ and the rest of $\boldsymbol{\pi}$ are fixed. Therefore, the necessary condition for

$G \sim G'$ becomes:

$$\mathbb{P}(x_{\pi_i}|\mathbf{x}(\mathcal{P}_i))\mathbb{P}(x_{\pi_j}|\mathbf{x}(\mathcal{P}_j)) \prod_{k\in\mathcal{C}_i\cup\mathcal{C}_j} \mathbb{P}(x_{\pi_k}|\mathbf{x}(\mathcal{P}_k)) = \mathbb{P}(x_{\pi_j}|\mathbf{x}(\mathcal{P}_i))\mathbb{P}(x_{\pi_i}|\mathbf{x}(\mathcal{P}_j)) \prod_{k\in\mathcal{C}_i\cup\mathcal{C}_j} \mathbb{P}(x_{\pi_k}|\mathbf{x}(\mathcal{P}'_k)) \tag{8}$$

**Proof of the first direction.** We first show that if $\mathcal{P}_i = \mathcal{P}_j$ and $\mathcal{C}_i = \mathcal{C}_j$ then condition (8) holds. If $\mathcal{P}_i = \mathcal{P}_j$ then $\mathbb{P}(x_{\pi'_i}|\mathbf{x}(\mathcal{P}'_i)) = \mathbb{P}(x_{\pi_j}|\mathbf{x}(\mathcal{P}_i)) = \mathbb{P}(x_{\pi_j}|\mathbf{x}(\mathcal{P}_j))$ and similarly $\mathbb{P}(x_{\pi'_j}|\mathbf{x}(\mathcal{P}'_j)) = \mathbb{P}(x_{\pi_i}|\mathbf{x}(\mathcal{P}_j)) = \mathbb{P}(x_{\pi_i}|\mathbf{x}(\mathcal{P}_i))$, which shows that the product of the first two CPDs in both graphs are equal.

Next, if $\mathcal{C}_i = \mathcal{C}_j$ then $\mathcal{C}_i \cup \mathcal{C}_j = \mathcal{C}_i \cap \mathcal{C}_j$. This means that $k$ can only be the child of both $i$ and $j$ (not just single one of them). Therefore $\mathcal{P}'_k = \mathcal{P}_k$ which results in the equality of the third terms $\mathbb{P}(x_{\pi_k}|\mathbf{x}(\mathcal{P}_k)) = \mathbb{P}(x_{\pi_k}|\mathbf{x}(\mathcal{P}'_k))$, which completes the proof of the first direction.

**Proof of the second direction.** We prove the contrapositive. For that, assume $\mathcal{P}_i \neq \mathcal{P}_j$ or $\mathcal{C}_i \neq \mathcal{C}_j$, then we should show that there exist $\mathbf{x}$ and $\boldsymbol{\theta}$ for which condition (8) gets violated. Below are the two branches of the proof by contrapositive:

- $\mathcal{P}_i \neq \mathcal{P}_j$: Consider $x_{\pi_i} = x_{\pi_j} = 0$, $\exists l \in \mathcal{P}_i - \mathcal{P}_j$ s.t. $x_l = 1$, $\forall l \in \mathcal{P}_j : x_l = 0$, and the rest of random variables can have arbitrary values. In words, $i$ and $j$ are inactive, all of $j$'s parents are inactive, and there exists at least one parent of $i$ which is active. For this setup, $\forall k \in \mathcal{C}_i \cup \mathcal{C}_j : \mathbb{P}(x_{\pi_k}|\mathbf{x}(\mathcal{P}_k)) = \mathbb{P}(x_{\pi_k}|\mathbf{x}(\mathcal{P}'_k))$, i.e., descendants of $i$ and $j$ are untouched.

  We show that there exist $\boldsymbol{\theta}$ for which the product of the terms 1 and 2 can not be equal. For the basic DBN and measurement error models, in $G$ we have $\mathbb{P}(x_{\pi_i} = 0|\mathbf{x}(\mathcal{P}_i))\mathbb{P}(x_{\pi_j} = 0|\mathbf{x}(\mathcal{P}_j)) = (1-\theta_{\pi_i}) \times 1$ and in $G'$ the product will be $\mathbb{P}(x_{\pi_j} = 0|\mathbf{x}(\mathcal{P}_i))\mathbb{P}(x_{\pi_i} = 0|\mathbf{x}(\mathcal{P}_j)) = (1-\theta_{\pi_j}) \times 1$. So for any $\theta_{\pi_i} \neq \theta_{\pi_j}$ the likelihoods are different. In other words, unless $\theta_{\pi_i} = \theta_{\pi_j}$ we have $\mathbb{P}(\mathbf{x}; G, \boldsymbol{\theta}) \neq \mathbb{P}(\mathbf{x}; G', \boldsymbol{\theta})$, which completes the proof for this case. Similarly, for the spontaneous activation model the condition reduces to $(1-\theta_{\pi_i})(1-\varepsilon_{\pi_j}) = (1-\theta_{\pi_j})(1-\varepsilon_{\pi_i})$, which can be violated in many ways, e.g., when $\theta_{\pi_i} = 1$ and while the rest of parameters are less than one.

- $\mathcal{C}_i \neq \mathcal{C}_j$: Above we showed that if $G \sim G'$ then $\mathcal{P}_i = \mathcal{P}_j$, therefore the product of the first two terms are equal in both graphs. So, here we only need to show that if $\mathcal{C}_i \neq \mathcal{C}_j$ the third terms $(\prod_{k\in\mathcal{C}_i\cup\mathcal{C}_j} \mathbb{P}(x_{\pi_k}|\mathbf{x}(\mathcal{P}_k)))$ can be different in both graphs. In this case, condition (8) reduces to the following necessary condition for $G \sim G'$:

$$\begin{aligned}&\prod_{k\in\mathcal{C}_i\cap\mathcal{C}_j} \mathbb{P}(x_{\pi_k}|\mathbf{x}(\mathcal{P}_k)) \prod_{l\in\mathcal{C}_i-\mathcal{C}_j} \mathbb{P}(x_{\pi_l}|\mathbf{x}(\mathcal{P}_l)) \prod_{k\in\mathcal{C}_j-\mathcal{C}_i} \mathbb{P}(x_{\pi_k}|\mathbf{x}(\mathcal{P}_k))\\ &=\\ &\prod_{k\in\mathcal{C}_i\cap\mathcal{C}_j} \mathbb{P}(x_{\pi_k}|\mathbf{x}(\mathcal{P}'_k)) \prod_{l\in\mathcal{C}_i-\mathcal{C}_j} \mathbb{P}(x_{\pi_l}|\mathbf{x}(\mathcal{P}'_l)) \prod_{k\in\mathcal{C}_j-\mathcal{C}_i} \mathbb{P}(x_{\pi_k}|\mathbf{x}(\mathcal{P}'_k))\end{aligned} \tag{9}$$

  For the shared children of $i$ and $j$ ($k \in \mathcal{C}_i \cap \mathcal{C}_j$) $\mathbb{P}(x_{\pi_k}|\mathbf{x}(\mathcal{P}_k)) = \mathbb{P}(x_{\pi_k}|\mathbf{x}(\mathcal{P}'_k))$ because $\mathcal{P}'_k = \mathcal{P}_k$, which leaves us with the latter two products in (9). Consider $x_{\pi_i} = 1$, $x_{\pi_j} = 0$, $\forall k \in \mathcal{C}_j - \mathcal{C}_i : \mathbf{x}(\mathcal{P}_k) = \mathbf{0}, x_{\pi_k} = 0$, and $\forall l \in \mathcal{C}_i - \mathcal{C}_j : \mathbf{x}(\mathcal{P}_l) = \mathbf{0}, x_{\pi_l} = 0$. The rest of random variables can have arbitrary values. Then, the above condition for basic DBN and measurement error models gets instantiated as:

$$\prod_{l\in\mathcal{C}_i-\mathcal{C}_j} (1-\theta_{\pi_l}) = \prod_{k\in\mathcal{C}_j-\mathcal{C}_i} (1-\theta_{\pi_k}),$$

and for spontaneous activation model as:

$$\prod_{l \in \mathcal{C}_i - \mathcal{C}_j} (1 - \theta_{\pi_l}) \prod_{k \in \mathcal{C}_j - \mathcal{C}_i} (1 - \varepsilon_{\pi_k}) = \prod_{k \in \mathcal{C}_j - \mathcal{C}_i} (1 - \theta_{\pi_k}) \prod_{l \in \mathcal{C}_i - \mathcal{C}_j} (1 - \varepsilon_{\pi_l}).$$

Clearly, one can devise parameters ($\theta$s and $\varepsilon$s) that violate above conditions. For example, with single $\theta_{\pi_l} = 1$ and the rest of parameters in $(0, 1)$, the LHS will be zero and the RHS non-zero, which violates both conditions. This example shows that if $\mathcal{C}_i \neq \mathcal{C}_j$ then $\exists \mathbf{x}, \boldsymbol{\theta}$ s.t. $\mathbb{P}(\mathbf{x}; G, \boldsymbol{\theta}) \neq \mathbb{P}(\mathbf{x}; G', \boldsymbol{\theta})$, which completes the proof.

■

*D.3. Canonical Forms*

Here, we define the canonical form of a given graph as a graph resulted from removing all redundant edges and uniquely labeling similar vertices.

**Definition 6.** Let $G$ be a DAG and let $(\mathbf{O}, \boldsymbol{\pi})$ be the decomposition of $G$ into an upper triangular matrix and a permutation. The **canonical form** of $G$ is the graph that is equivalent to $G$ with no redundant edges along with a permutation $\boldsymbol{\pi}$ such that each set of similar nodes is ordered from least index to greatest index.

The canonical form of $G$ is in fact a canonical form– it defines a unique DAG, which helps us recovering a unique graph from all members of an equivalent class. In practice, this property will helps a lot in visualizing the output and understanding the recovered BN structure.

**Theorem 7.** *$G \sim G'$ if and only if $G$ and $G'$ have the same canonical form.*

*Proof.* The proof is the direct application of previous propositions (Propositions 1, 2, and 3) for redundant edges and similar vertices. ■

**References**

Daphne Koller and Nir Friedman. *Probabilistic graphical models: principles and techniques.* MIT press, 2009.

I M Oliver, D J Smith, and J R C Holland. A study of permutation crossover operators on the traveling salesman problem. In *Proc. 2nd Int. Conf. Genetic Alg.*, pages 224–230, Hillsdale, NJ, USA, 1987. L. Erlbaum Associates Inc.

Aniko Szabo and Kenneth Boucher. Estimating an oncogenetic tree when false negatives and positives are present. *Math. Biosci.*, 176(2):219–236, April 2002.
